# Polygyny carries costs in both sexes in Trinidadian guppies

**DOI:** 10.64898/2026.04.07.716995

**Authors:** Tess Makenna van der Walle, Flavio Di Giorgio, Tomos Potter, Anja Felmy

## Abstract

According to sexual selection theory, males should benefit more from mating with multiple partners than females do, as male investment into offspring production is typically lower. For females, empirical evidence indeed often shows diminishing returns or even costs of mating multiply. For males, the assumption often seems to be “the more, the better” – i.e., a steady increase of male reproductive success with mate number – but experimental tests of it are rare. Here we used a laboratory experiment with Trinidadian guppies (*Poecilia reticulata*), known for being promiscuous, to assess how pairing males weekly with 4 vs. 7 females affects both sexes’ reproductive performance (n = 32 polygynous males and 170 monogamous females). Increased polygyny delayed females’ reproductive onset by 9% and tripled their risk of reproductive failure. High-polygyny males fathered offspring with 49% more females and had 73% higher daily reproductive output. Yet, they needed 19% longer to initiate pregnancy, and only accumulated more offspring than low-polygyny males after two months. This study suggests that male mating performance is not unlimited. Especially when high extrinsic mortality selects for fast reproduction, less polygyny might be advantageous, and the strength of sexual selection perhaps more similar between the sexes than often assumed.

## Introduction

Sexual selection theory posits that males – i.e. the sex with the smaller, more numerous gametes – benefit more from mating with additional partners than females do [1]. The slope of reproductive success on mate number, called the “Bateman gradient”, is thus expected to be steeper in males, and sexual selection consequently stronger [2-4]. Across the animal kingdom, empirical evidence indeed supports these expectations [5].

The costs and benefits of multiple mating are relatively well-studied in females. Polyandry (i.e. female matings with multiple males) has been found to be common in a wide range of taxonomic groups [e.g. 6, 7-9], despite diminishing returns or even costs as the number of mating partners increases [e.g. 10, 11-13]. By contrast, polygyny, where one male mates with multiple females, is generally assumed to benefit males at a cost to females [14, 15]. Even at high mating rates, males are expected to gain from additional copulations without saturation, because mating with more females should equal more offspring per male [2, 14, 15]. When costs of polygyny to males have been studied, it was mainly with respect to increased male-male competition [e.g. 16, 17] or reduced amounts of paternal care given to offspring [e.g. 18, 19, 20].

However, polygyny might be costly for males even when male-male competition and male parental investment are absent. Although sperm has long been viewed as an unlimited resource [2, 21], a male’s number of ejaculates is limited [22]. Empirical evidence of sperm depletion, such as reduced ejaculate size and fertilisation rates after successive matings, exists e.g. for Soay sheep [23], Medaka fish [24], and Lepidopterans [25]. Also ejaculate components other than sperm can become depleted, depressing female fecundity [26]. Even without sperm or ejaculate depletion, male mating performance can decrease over successive matings [27]. Hence, if males spread themselves too thinly by doing a bad job of mating with too many females, the costs to their females might translate into costs to themselves, and their reproductive success might be lower than that of less polygynous males.

Here we test whether polygyny imposes costs on male or female reproduction in Trinidadian guppies (*Poecilia reticulata*). Guppies are internally fertilising, live-bearing fish whose promiscuous mating system makes them an excellent system for studying sexual selection [28, 29]. Neither sex provides parental care, removing one of the primary costs of – and thus limiting factors to – polygyny. We conducted a laboratory experiment, pairing males weekly with 4 vs. 7 females in sequential monogamy. By assigning males a set level of polygyny, we remove the confounding effect of male quality that, in observational settings, can prevent lower-quality males from achieving higher degrees of polygyny, potentially masking its costs. We measured female age at first reproduction and first litter size and computed male and female daily and cumulative reproductive output. We found that increased polygyny was costly for both sexes. High-polygyny males were slower and less successful at initiating pregnancy, thereby delaying high-polygyny females’ reproductive onset and tripling their risk of reproductive failure.

## Methods

### Experimental fish

The fish used here were first-generation (F1) laboratory-born offspring of 297 founders collected from a high-predation stream (Guanapo River) in Trinidad, and brought to Lund University, Sweden, in October 2024. They were housed in a temperature-controlled room (26.5 ± 0.2ºC [mean ± SD]) in four independent recirculation systems with continuous filtration, ultraviolet sterilisation, and a 12L:12D light cycle. Founder females were kept in individual 1.1-litre tanks and checked daily for fry. F1 fry were reared in 1.1-litre tanks in sibling groups (maximum ten fry per tank). At age 30-35 days, F1 juveniles were anaesthetised, sexed, and moved to individual 1.1-litre tanks. All fish were fed *ad libitum* with liver paste and live brine shrimp nauplii (*Artemia* spp.).

### Mating trials

From the sexed F1 juveniles, we selected 170 females and 32 males. They had 75 founder mothers, with 2.3 ± 1.0 F1 females per founder mother (range 1-6) and a different founder mother for every F1 male. We assigned fish to one of two experimental levels of polygyny. At the low level, a male was paired with four females (n = 18 males, n = 18 x 4 = 72 females), at the high level, with seven females (n = 14 males, n = 14 x 7 = 98 females). We thus had 32 mating groups, each consisting of one male and either four or seven females, respectively.

Males had different founder mothers than their females. One individual initially presumed female was later found to be male, and one female died early, reducing the number of females in two high-polygyny groups to six, and the total female number to 168.

Within each mating group, we paired fish according to a two-week schedule, such that each female was mated once a week, and males had regular recovery periods (Fig. 1). The schedule alternated both the females’ mating order and day-night cycle of mating opportunities, to equalise the likelihood of successful insemination. We set up mating groups in a staggered fashion over 37 days (Supporting table 1).

**Figure 1.**
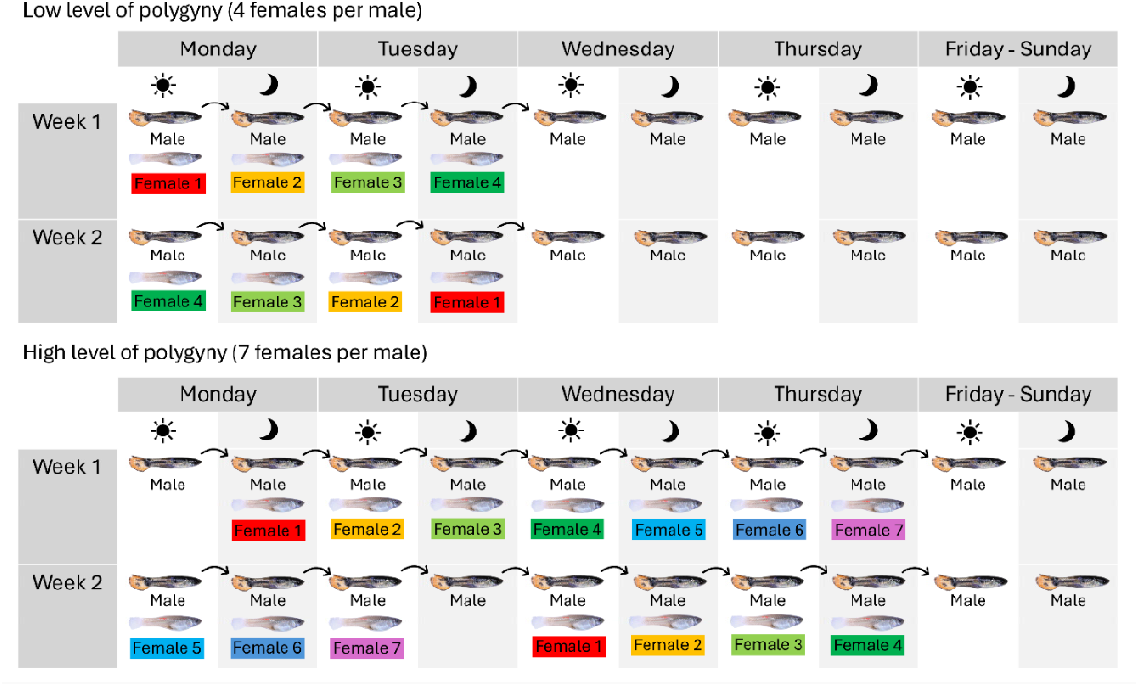
Schedule of mating trials at two experimental levels of polygyny. Mating groups contained one male and either four females (low polygyny, n = 18 groups) or seven females (high polygyny, n = 14 groups). All fish were initially virgins. Females were paired with their assigned male once a week, either during daytime (approx. 8 hours) or overnight (approx. 16 hours). The schedule was repeated until females either gave birth or reached 100 days of age. Females were housed in individual tanks; males were moved between female tanks, with recovery periods in their own tank. Hence, matings happened in a sequentially monogamous way and always in the females’ home tank.

Females were mated first at age 42.7 ± 4.8 days (range 34-64) and males at age 51.3 ± 7.1 days (range 21-59). We ensured that all males were sexually mature, as indicated by a fully developed gonopodium. Low-polygyny females were, on average, 4.2 days younger on their first mating opportunity than high-polygyny females. In part, this difference (*t* = -6.2, *df* = 154.8, *p* = 6.3e-09) arose because fewer days are needed to mate one male with four rather than seven females (see Fig. 1), but logistic difficulties contributed to it. To account for a potential bias introduced by variation in female age at first mating, we included it as a predictor in statistical models.

We mated females until they gave birth or reached 100 days of age without reproducing. In four mating groups, the male died before all females either reproduced or turned 100 days old. The resultant lack of mating opportunities for these 16 females (8 from high-, 8 from low-polygyny groups) could affect their reproductive performance. “Death of male” was thus a predictor in statistical models. Non-reproductive females aged 100 days were kept isolated for 12 days. If by day 112 they still had not given birth, we declared them non-reproductive with respect to their assigned male. To investigate whether they were infertile, we then paired them with two additional males. These were unrelated to the females, the females’ first males, and each other. None of them had previously been part of mating trials. Every week, we paired non-reproductive females with their two males, sequentially, for 24 hours per male. Females that were non-reproductive by the age of 165 days were considered infertile.

### Measurement of phenotypic traits

We measured female age at first parturition and the number of offspring in first litters. Female tanks were checked daily for fry. Fry born within one day of each other were considered to be part of the same litter. We calculated female daily reproductive output by dividing first litter size by the interval between a female’s first mating opportunity and first parturition. A male’s daily reproductive output was the sum of the daily reproductive outputs of his assigned females. We identified the mating opportunity a female’s pregnancy was likely initiated at by subtracting the shortest observed gestation length (22 days) from her first parturition date, followed by finding the closest mating opportunity before that date.

### Statistical analysis

We used generalised linear mixed models (GLMMs) to analyse the interval between females’ first mating opportunity and first birth, the size of the first litter, and female daily reproductive output. The fixed effects were level of polygyny (factor: low vs. high), female age at first mating, female pairing order in week 1 of mating trials, premature death of male (factor: no vs. yes), and the recirculation system a female was housed in (factor: 1-4). We fitted random intercepts for male identity and female’s mother’s identity, to account for non-independence of data among females grouped by these factors.

We compared models fitted with different error structure to find the best-fitting one for each trait. For the mating-to-birth interval, the best model had Gaussian errors, an identity link function, and base-*e* log-transformed response values; for first litter size, Poisson errors, a log link, and untransformed response values; for female daily reproductive output, Gaussian errors, an identity link, and a square-root-transformed response.

We performed statistical analyses in R v. 4.4.2 [30]. GLMMs were fitted using “glmmTMB” [31]. We tested the significance of random effects with log-likelihood ratio tests and assessed model fit using DHARMa [32]. Scatterplots were prepared using “beeswarm” [33]. Values are given as mean ± SD.

## Results

### Effects of polygyny on female reproduction

Increased polygyny delayed females’ reproductive onset (Fig. 2a, Table 1). Females in the high-polygyny treatment had 9.0% longer intervals between their first mating opportunity and first birth (42.9 ± 11.4 days) than those in the low-polygyny treatment (39.4 ± 8.8 days).

**Figure 2.**
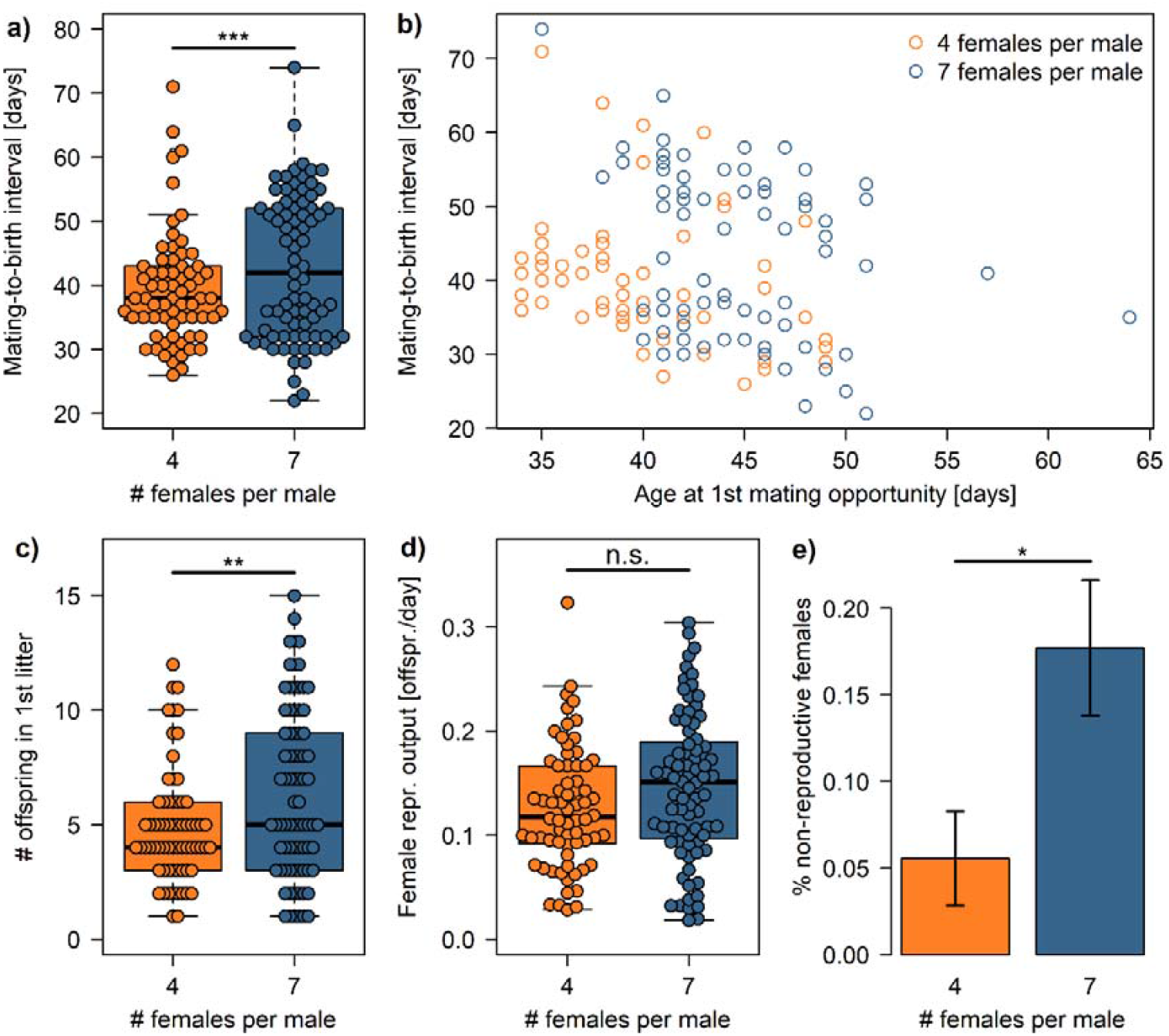
Effects of experimentally increased polygyny on female reproductive performance. High-polygyny females needed more time to conceive than low-polygyny females (a) – an interval that also increased the younger a female was when mated for the first time (b). High-polygyny females had larger first litters (c) but equal daily reproductive output (d). They were, however, more likely to remain non-reproductive (e). In (e) the sample size is 72 and 96 females per group, respectively, and error bars are binomial standard errors (± 1 SE) computed according to Zar [34].

**Table 1.** Outcome of GLMMs for reproductive traits of female guppies whose partners experienced either a lower or higher level of polygyny. Number of females = 147 (lower level of polygyny: 68, higher level of polygyny: 79).

|  | Mating-to-birth interval |  |  | No. offspring in first litter |  |  | Daily reproductive output |  |  |
| --- | --- | --- | --- | --- | --- | --- | --- | --- | --- |
| **Fixed effects:** | **Estimate**<br>**(S.E.)** | **z-value** | **p-value** | **Estimate**<br>**(S.E.)** | **z-value** | **p-value** | **Estimate**<br>**(S.E.)** | **z-value** | **p-value** |
| (Intercept) | 4.34 (0.24) | 18.00 | < 2e-16 | 1.59 (0.46) | 3.44 | 0.0006 | 0.28 (0.07) | 3.78 | 0.0002 |
| No. female partners (4 vs. 7) | 0.22 (0.06) | 3.53 | 0.0004 | 0.30 (0.11) | 2.61 | 0.0091 | 0.02 (0.02) | 0.91 | 0.36 |
| Female age at first mating | -0.02 (0.01) | -2.78 | 0.0054 | 0.00 (0.01) | 0.04 | 0.97 | 0.00 (0.00) | 1.00 | 0.32 |
| Female pairing order | -0.02 (0.01) | -1.56 | 0.12 | -0.03 (0.03) | -1.13 | 0.26 | -0.00 (0.00) | -0.38 | 0.70 |
| Death of male (no vs. yes) | 0.20 (0.09) | 2.25 | 0.0242 | 0.18 (0.17) | 1.03 | 0.30 | 0.00 (0.03) | 0.01 | 0.99 |
| Recirculation system 2 | 0.08 (0.06) | 1.34 | 0.18 | -0.08 (0.14) | -0.54 | 0.59 | -0.03 (0.02) | -1.30 | 0.20 |
| Recirculation system 3 | -0.04 (0.06) | -0.73 | 0.47 | 0.02 (0.13) | 0.12 | 0.90 | 0.01 (0.02) | 0.32 | 0.75 |
| Recirculation system 4 | -0.00 (0.05) | -0.08 | 0.94 | 0.13 (0.13) | 1.02 | 0.31 | 0.01 (0.02) | 0.68 | 0.50 |
| **Random effects:** | **Var (S.D.)** | **$\chi^2_1$** | **p-value** | **Var (S.D.)** | **$\chi^2_1$** | **p-value** | **Var (S.D.)** | **$\chi^2_1$** | **p-value** |
| Male identity | 0.014 (0.119) | 13.41 | 0.0003 | 0.019 (0.139) | 1.72 | 0.19 | 0.000 (0.000) | 0.0 | 1.00 |
| Female's mother's identity | 0.007 (0.084) | 2.39 | 0.12 | 0.054 (0.233) | 7.23 | 0.0072 | 0.000 (0.000) | 0.0 | 1.00 |
| Residual | 0.030 (0.174) |  |  |  |  |  | 0.008 (0.091) |  |  |

Mating-to-birth intervals were also longer for younger females (Fig. 2b) and when a female’s assigned male died prematurely (Table 1). They did not depend on the pairing order of females, nor on their placing inside the laboratory (Table 1). Male identity explained a significant amount of variation in mating-to-birth intervals, while the identity of the females’ mother did not (Table 1).

High-polygyny females had larger first litters than low-polygyny females (Fig. 2c) but the same daily reproductive output (Fig. 2d, Table 1). On average, first litters contained 30.8% more offspring in high-polygyny groups (6.4 ± 3.7 vs. 4.9 ± 2.5; Fig. 2c). Given that litter size was positively correlated with female age at first parturition (Pearson’s *r* = 0.49, *t*_145_ = 6.8, *p* = 2.04e-10), this difference likely arose from high-polygyny females being older and thus presumably larger when producing their first litters. All other predictors of first litter size were nonsignificant, except for the females’ mothers’ identity (Table 1). However, despite their larger first litters, high-polygyny females did not have a higher daily rate of offspring production: 0.15 ± 0.07 vs. 0.13 ± 0.06 offspring per day (Fig. 2d, Table). No other predictor influenced that rate either (Table 1).

The high-polygyny treatment increased females’ risk of reproductive failure (Fig. 2e). Only 21 out of 168 females (12.5%) did not reproduce, but rates of non-reproduction were over three times greater in the high- (17.7%) compared to the low-polygyny group (5.6%; Pearson’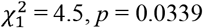 Fig. 2e). Notably, non-reproductive females were not infertile: upon being paired with additional males, 20 out of 21 females (95.2%) gave birth.

### Effects of polygyny on male reproduction

At first glance, being more polygynous boosted males’ reproductive success. Males in high-polygyny groups fathered offspring with 49.4% more females (5.6 ± 1.3) than low-polygyny males (3.8 ± 0.5; Wilcoxon rank sum test: *W* = 31, *p* = 9.33e-05; Fig. 3a). They also had a 73.0% higher total daily reproductive output (*W* = 38, *p* = 0.0005; Fig. 3b), receiving 0.83 ± 0.31 compared to 0.48 ± 0.14 offspring per day from all their females combined. This was despite a 3.2 times higher fraction of non-reproductive females among high-than low-polygyny males (17.5 ± 18.7% vs. 5.6 ± 13.7%; *W* = 71, *p* = 0.0161).

**Figure 3.**
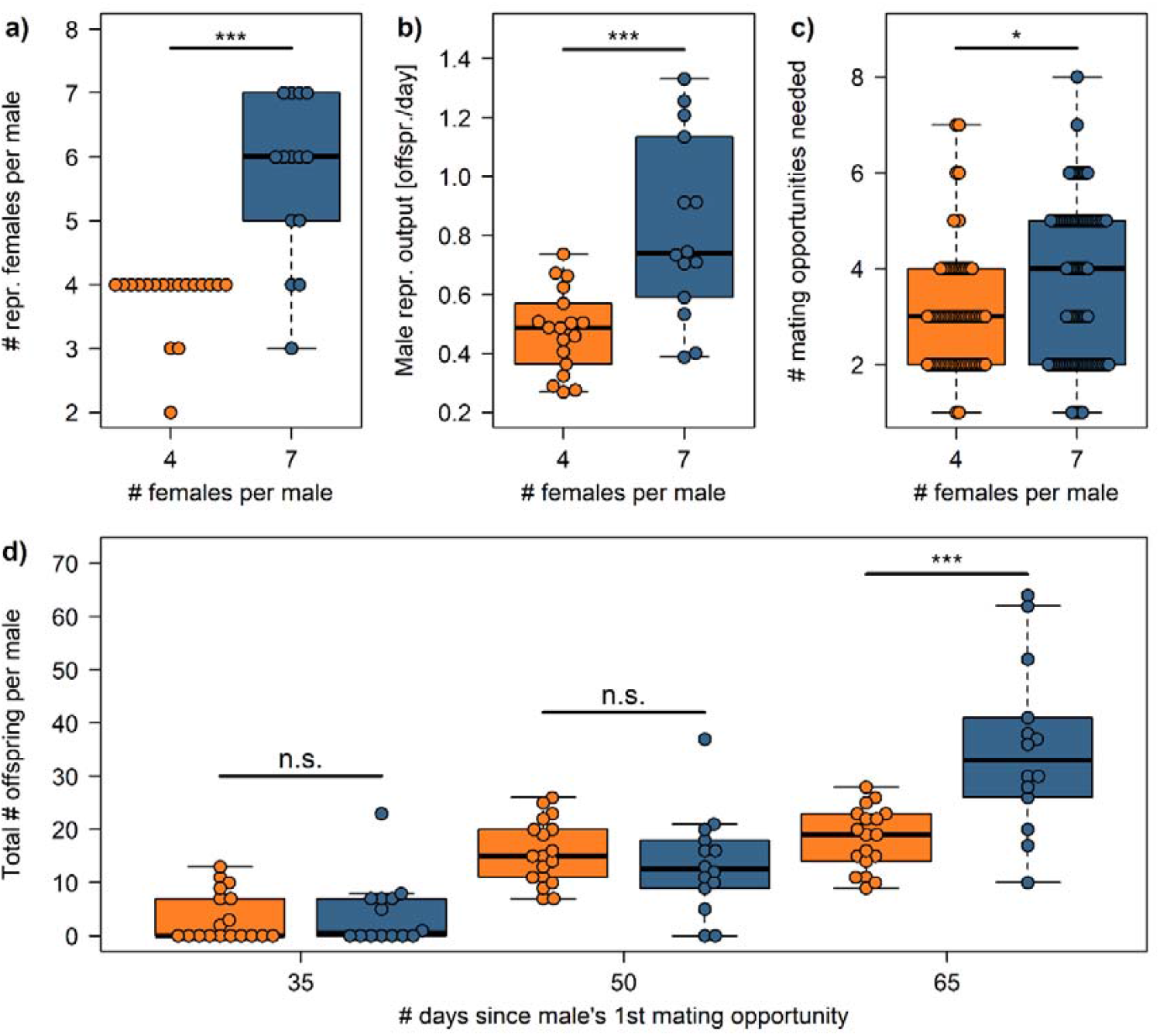
Effects of experimentally increased polygyny on male reproductive performance. High-polygyny males gained offspring from a higher absolute number of females (a) and had a higher daily reproductive output (b). However, they needed more mating opportunities to impregnate their females (c) and only accumulated more offspring than low-polygyny males 65 days after their first mating opportunity (d).van der Walle *et al*. – Manuscript

However, when taking the aspect of time into account, increased polygyny was costly for males. High-polygyny males needed 18.5% more mating opportunities – and thus 18.5% more time – to initiate pregnancy in a female (3.7 ± 1.7 vs. 3.1 ± 1.3 mating opportunities; Welch two-sample *t*-test: *t*_142.3_ = -2.4, *p* = 0.0201; Fig. 3c). Additionally, the benefits of increased polygyny accrued relatively late. The total number of offspring a high-polygyny male sired only surpassed that of low-polygyny males 65 days after a male’s first mating (*W* = 36, *p* = 0.0007); after 35 and 50 days, respectively, the difference was nonsignificant (both *p* > 0.34; Fig. 3d).

## Discussion

Our findings indicate that polygyny imposes direct costs on female reproduction, and that these translate into reproductive costs to males. Female guppies in the high-polygyny treatment conceived more slowly and were three times more likely to remain non-reproductive. High-polygyny males, for their part, were slower at initiating pregnancy, had higher reproductive failure, and only outperformed low-polygyny males after two months. While females in our study undoubtedly faced higher costs than males given their inability to seek compensatory matings elsewhere, we show that polygyny carries direct fitness costs for male guppies, too. We can rule out that high-polygyny males were accidentally paired with less fertile females, as non-reproductive females successfully reproduced with other males.

Costs of polygyny are rarely considered in species without parental care. We are not aware of prior work in guppies that quantified how polygyny affects male or female reproduction. But sexual interactions have been found to be costly to males in other respects. Large guppy males show higher mortality when housed in mixed-sex compared to all-male groups [35]. Increased male reproductive effort decreases males’ foraging rates, growth, and survival [36, 37]. Sperm production is plastic in guppies, decreasing under food limitation [38] and increasing in the presence of females [39-41], suggesting that its costs are substantial (although see [42]). Data on how male mating history affects female reproductive performance are sparse; guppy females primarily experience polyandry [28, 29], where matings with multiple males likely mitigate some of the costs associated with polygyny.

The delayed conception and first birth of high-polygyny females suggest that polygyny might be costly especially when extrinsic mortality is high. When mortality risks are high, anything that lowers a male’s ability to fertilise eggs quickly should be selected against. After all, he might die before encountering her again, or she might die before conceiving or before giving birth. Guppies inhabiting downstream river sections, where population densities are kept low by intense predation [43, 44], face substantially higher monthly mortality rates than their conspecifics in upstream sections [45]. It might thus be expected that the optimum degree of polygyny is lower in high-than in low-predation populations – thus adding another facet to the many ways in which sexual selection dynamics differ between high- and low-predation guppies [46-49]. Also irrespective of extrinsic mortality it might be advantageous to reproduce early. There is evidence across various species that earlier-born offspring have higher fitness than later-born offspring, for example in lizards [50], roe deer [51], and mosquitofish [52]. Moreover, in a growing population, females gain more fitness from offspring born earlier, as reproducing earlier allows them to contribute to population growth sooner [53-55].

Several limitations of our study should be addressed. First, it cannot pinpoint the mechanism behind the delayed reproductive onset of high-polygyny females, which might be mating fatigue, sperm limitation, or both. Future work could address this, perhaps by observing mating behaviour or measuring sperm counts. Second, our experimental design simplified natural conditions in three important ways: by imposing monogamy on females, considering first litters only, and eliminating male-male competition. In the following, we explain why we believe these simplifications do not interfere with our study’s goal and instead make our estimated costs of polygyny conservative. Females being monogamous is an unlikely scenario; in the field, both courtship displays and sneak mating attempts are common [28] and levels of multiple paternity typically high [29]. Consequently, females that failed to get pregnant quickly from their high-polygyny male would likely be inseminated by other males, thereby increasing male costs of polygyny. The restriction to first litters might, by contrast, not be overly unrealistic: under natural conditions, females might not live to produce a second litter or might sire it with sperm from another male. Nonetheless, had our study included second litters, then many low-polygyny females would have produced second litters before high-polygyny females had their first, reducing the benefit of increased polygyny even after 65 days. Finally, eliminating male-male competition allowed us to quantify effects of polygyny independent of variation in males’ competitive ability before or after copulation. This, again, makes our study conservative: if a more polygynous mating strategy does not pay off in a benign, competition-free scenario, it might not under more realistic conditions either. Future experiments or modelling work could address costs of polygyny in the context of competition between males.

An unexpected finding is the apparent bimodal distribution of the mating-to-birth interval, visible as two parallel, downward-facing clouds in Fig. 2b. In the lower cloud, mating-to-birth intervals were relatively short (~22-48 days); in the upper cloud, relatively long (~40-75 days). Within clouds, interval length depended on female age at first mating, decreasing the closer that age was to about 50 days, when females apparently reached sexual maturity. Given that guppy females, unlike males, lack external signs of maturity [56], identifying that age in females is an interesting result in and of itself. If a female’s first mating happened around age 50 days and was successful, the mating-to-birth interval was shortest (min. 22 days), consisting only of the gestation time. However, if the mating was (supposedly) unsuccessful, the interval increased to about twice that length (~42-53 days). At this point, we do not know whether this bimodality is an artefact of pairing females discontinuously (i.e. once a week); more data will need to establish its veracity. However, it is faintly reminiscent of a spontaneous ovarian cycle, for which circumstantial evidence exists in other poeciliids [57].

In conclusion, our study shows that costs of polygyny are borne by females as well as males. This implies that sexual selection operates on females, too – if polygyny comes with reproductive costs in males, then presumably there is male mate choice. The appreciation of sexual selection being important in females is growing, mainly driven by widespread positive associations between polyandry and female reproductive success [58, 59]. We contribute to this paradigm shift by highlighting that male mating performance is not unlimited, and the association between polygyny and male reproductive success perhaps not unconditionally positive.

## Ethics

This research was approved by the Ethical Committee on Animal Experiments, Lund/Malmö, Sweden (Dnr 5.8.18-13832/2024).

## Data accessibility

All data and R code associated with this manuscript will be available as part of the Supporting Information following manuscript acceptance.

## Declaration of AI use

We have not used AI-assisted technologies in creating this article.

## Author contributions

T.V.D.W.: investigation, formal analysis, writing – original draft.

F.D.G.: investigation, writing – review & editing, supervision.

T.P.: formal analysis, writing – review & editing.

A.F.: conceptualisation, formal analysis, data curation, writing – original draft, writing – review & editing, visualisation, supervision, funding acquisition.

## Acknowledgements

We thank the core scientists of The Guppy Project, David Reznick, Joe Travis, Ron Bassar, and Tim Coulson, for letting us stay in their field station in Trinidad and using its infrastructure for capturing the founders of our laboratory population. We especially thank Ryan Mohammed for his invaluable help with export permits and logistics, and Ignacio Paulin and Gabriela Jeliazkov for help in the field. In Lund, we thank Madalena Madeira for help with fish maintenance, and Olof Berglund for comments on an earlier version of the manuscript.

