## Supporting Table 1 for "Polygyny carries costs in both sexes in Trinidadian guppies"

| Mating group | No. females per male | First day of pairing | Mating group | No. females per male | First day of pairing |
|---|---|---|---|---|---|
| 1 | 7 | 09.12.2024 | 15 | 4 | 30.12.2024 |
| 2 | 7 | 09.12.2024 | 16 | 4 | 30.12.2024 |
| 3 | 7 | 16.12.2024 | 17 | 4 | 30.12.2024 |
| 4 | 7 | 16.12.2024 | 18 | 4 | 30.12.2024 |
| 5 | 7 | 16.12.2024 | 19 | 4 | 30.12.2024 |
| 6 | 7 | 23.12.2024 | 20 | 4 | 30.12.2024 |
| 7 | 7 | 23.12.2024 | 21 | 4 | 06.01.2025 |
| 8 | 7 | 23.12.2024 | 22 | 4 | 06.01.2025 |
| 9 | 7 | 23.12.2024 | 23 | 4 | 06.01.2025 |
| 10 | 7 | 23.12.2024 | 24 | 4 | 06.01.2025 |
| 11 | 7 | 23.12.2024 | 25 | 4 | 06.01.2025 |
| 12 | 7 | 23.12.2024 | 26 | 4 | 06.01.2025 |
| 13 | 7 | 23.12.2024 | 27 | 4 | 06.01.2025 |
| 14 | 7 | 23.12.2024 | 28 | 4 | 06.01.2025 |
| | | | 29 | 4 | 06.01.2025 |
| | | | 30 | 4 | 06.01.2025 |
| | | | 31 | 4 | 15.01.2025 |
| | | | 32 | 4 | 15.01.2025 |

**Supporting Table 1. Date of first mating opportunity for each mating group.** On that date, the male was paired with his first assigned female. All high-polygyny groups (7 females per male), and six low-polygyny groups (4 females per male), were started in December, the others in early January. For high-polygyny groups, we adjusted the mating schedule for two weeks over the Christmas period, with the male moving from one female tank to the next every 24 hours, without recovery periods. No adjustments were made for low-polygyny groups.
